# HaploJuice: Accurate haplotype assembly from a pool of sequences with known relative concentrations

**DOI:** 10.1101/307025

**Authors:** Thomas K. F. Wong, Louis Ranjard, Yu Lin, Allen G. Rodrigo

**Author notes:** ^*^.

## Abstract

Pooling techniques, where multiple sub-samples are mixed in a single sample, are widely used to take full advantage of high-throughput DNA sequencing. Recently, Ranjard et al. [1] proposed a pooling strategy without the use of barcodes. Three sub-samples were mixed in different known proportions (i.e. 62.5%, 25% and 12.5%), and a method was developed to use these proportions to reconstruct the three haplotypes effectively. HaploJuice provides an alternative haplotype reconstruction algorithm for Ranjard et al.’s pooling strategy. HaploJuice significantly increases the accuracy by first identifying the empirical proportions of the three mixed sub-samples and then assembling the haplotypes using a dynamic programming approach. HaploJuice was evaluated against five different assembly algorithms, Hmmfreq [1], ShoRAH [2], SAVAGE [3], PredictHaplo [4] and QuRe [5]. Using simulated and real data sets, HaploJuice reconstructed the true sequences with the highest coverage and the lowest error rate. HaploJuice achieves high accuracy in haplotype reconstruction, making Ranjard et al.’s pooling strategy more efficient, feasible, and applicable, with the benefit of reducing the sequencing cost.

## Introduction

With the rapid advancement of next-generation sequencing technologies, it is possible to obtain several gigabases of sequences in a single day. Given the huge volume of throughput, it is often cost-effective to mix multiple sub-samples in a single sample for sequencing, a process called pooling. Several approaches have been developed to demultiplex the sequencing reads from the mixture, i.e. assign reads to their respective sub-samples. For example, a short unique identifiable sequence tag (i.e. barcode) is often appended to each DNA molecule of the same sub-sample before pooling and sequencing. Subsequently, the reads can be assigned to the appropriate sub-sample in the mixture based on their associated barcodes [6]. However, the cost of the sequencing library preparation increases linearly with the number of required barcodes.

Recently, Ranjard, et al. [1] proposed a pooling strategy that does not require the use of barcodes. The strategy consists in mixing sub-samples in different known proportions. These proportions induce different expected frequencies of the sub-sample variants in the mixture, and hence, different expected sequencing read coverages. These frequencies, in turn, allow the sub-sample sequences to be reconstructed accurately. Ranjard et al. applied their method to mitochondrial sequences from three kangaroo subsamples mixed in proportions 62.5%, 25%, and 12.5%, and showed that the three haplotypes could be assembled effectively, thus reducing the cost of sequencing significantly.

Ranjard et al. developed Hmmfreq [1] which requires alignment of short-reads to a suitable reference sequence. By assuming the sub-sample proportions 62.5%, 25%, and 12.5%, Hmmfreq uses a Dirichlet-multinomial model [7] and a Hidden Markov Model (HMM) to reconstruct the three haplotypes from the mixture of reads. Other methods [2–5] that also use reference sequences to align short reads, have been developed to recover sequences in mixtures, where the number of sub-samples and the corresponding proportions in the mixtures are unknown. For instance, ShoRAH [2] implements local-window clustering to recover the constituent sequences (or “haplotypes”) in a mixture. SAVAGE [3] uses an overlap graph and clique enumeration to reconstruct multiple haplotypes. PredictHaplo [4] uses Dirichlet prior mixture model, starts local reconstruction at the region of maximum coverage and progressively increases the region size until it covers the entire length of haplotypes. QuRe [5] uses sliding windows and reconstructs the haplotypes based on multinomial distribution matching heuristic algorithm [8]

**Table 1.** The results of estimation on the sample proportions by HaploJuice. 100 data sets were simulated for each case.

| Case | Actual sample proportion | | | Expected sample proportion<br>(Average ± Standard deviation) | | |
| --- | --- | --- | --- | --- | --- | --- |
| | $f_1$ | $f_2$ | $f_3$ | | | |
| 1 | 0.5 | 0.4 | 0.1 | $0.50 \pm 0.001$ | $0.40 \pm 0.001$ | $0.10 \pm 0.001$ |
| 2 | 0.5 | 0.3 | 0.2 | $0.50 \pm 0.001$ | $0.30 \pm 0.001$ | $0.20 \pm 0.001$ |
| 3 | 0.6 | 0.3 | 0.1 | $0.60 \pm 0.001$ | $0.30 \pm 0.001$ | $0.10 \pm 0.001$ |
| 4 | 0.7 | 0.2 | 0.1 | $0.70 \pm 0.001$ | $0.20 \pm 0.001$ | $0.10 \pm 0.001$ |

In this paper, we focus on the pooling strategy [1] proposed by Ranjard et al. but our method, however, does not assume any prior knowledge on the sample proportions; instead, we assume that the number of sub-samples in the mixture is known. We compute the sub-sample proportions directly from the mixture of reads. Based on the estimated sample proportions, we use multinomial model and dynamic programming to construct multiple haplotypes simultaneously. HaploJuice, which is an extension of Hmmfreq [1], considers all possible combinations for assigning local sub-sequences to haplotypes, and selects the combination with the highest overall likelihood. We evaluate HaploJuice against five different assembly algorithms, Hmmfreq [1], ShoRAH [2], SAVAGE [3], PredictHaplo [4] and QuRe [5], using simulated and real data sets in which three sequences are mixed in known frequencies. Based on our results, HaploJuice reconstructs sequences with the highest coverage of the true sequences and had the lowest error rate.

## Experimental results

HaploJuice first identifies the underlying sub-sample proportions from a mixture of reads and, second, reconstructs the haplotypes using these estimated proportions. As with Hmmfreq it requires an alignment of short-read sequences against a reference sequence. In our analysis, all reads are aligned to the reference sequence using Bowtie 2 [9].

Simulated datasets were used to evaluate our methods. Four hundred data sets were simulated and each data set was a mixture of three sub-samples. The three sub-samples were mixed under various proportions: 5:4:1, 5:3:2, 6:3:1, and 7:2:1 (100 data sets each). 150-long pair-ended reads with total coverage 1500x were simulated by ART [10] with the default Illumina error model from three 10k-long haplotypes, which were generated by INDELible [11] using JC [12] model from a 3-tipped tree with 0.05 root-to-tip distance randomly created by Evolver [13] from PAML [14] package.

After using Bowtie 2 [9] to align the reads against the root sequence (also reported from INDELible [11]), we ran HaploJuice to estimate the sub-sample proportions in the mixture. As shown in Table 1, on average, the estimated sub-sample proportions were the same as the actual proportions with standard deviation 0.001. The method of estimation on the sub-sample proportions is, therefore, found to be effective on these simulated data sets.

HaploJuice was then used to reconstruct the haplotype sequences for each data set based on the estimated sample proportions. HaploJuice was compared to five different assembly algorithms, including Hmmfreq [1], ShoRAH [2], SAVAGE [3], PredictHaplo [4] and QuRe [5]. Note that SAVAGE, PredictHaplo and QuRe do not have prior assumptions on the number of haplotypes, whereas HaploJuice and Hmmfreq do. MetaQUAST [15] was then used with default parameters to evaluate the contigs, which were resulted by all the software, against the true sequences. Table 2 shows the summary of the performance of different methods on the simulated data sets. On average, HaploJuice reconstructed contigs over 99.7% haplotype coverage, which was the highest among all the methods. When checking the error rates (i.e. the percentage of bases in the contig sequences having mutations or indels when compared against with the real haplotypes), HaploJuice was less than 0.005% on average. It was the lowest among the software which reconstructed contigs over 90% haplotype coverage. In conclusion, HaploJuice is shown effective from the simulated data sets.

**Table 2.** Comparison of performance of different methods on construction of three haplotypes for simulated data sets. 100 data sets were generated for each of the cases with different sets of sample proportions. Format of the data is: average ± standard deviation. The best value for each column is **highlighted** the software outputting the contigs over 90% haplotype coverage.

| a. Proportion of three samples: 0.5, 0.4, 0.1 (total length of three haplotypes: 30k) |  |  |  |  |  |
| --- | --- | --- | --- | --- | --- |
| Software | # contigs<br>$\geq 500\text{bp}$ | Longest<br>contig | N50 | Haplotypes<br>coverage % | Error rate % |
| HaploJuice | **3.0 ± 0.0** | **9975 ± 6.8** | **9971 ± 6.5** | **99.7 ± 0.0** | **0.001 ± 0.004** |
| hmmfreq [1] | **3.0 ± 0.0** | 9855 ± 6.8 | 9850 ± 6.3 | 98.5 ± 0.0 | 0.276 ± 0.254 |
| shoRAH [2] | 30.8 ± 11.7 | 9819 ± 124.8 | 9799 ± 116.7 | 97.5 ± 3.5 | 0.646 ± 0.492 |
| SAVAGE [3] | 9.8 ± 3.5 | 9972 ± 11.8 | 305 ± 300.3 | 51.3 ± 7.1 | 0.001 ± 0.004 |
| PredictHaplo [4] | 2.0 ± 0.2 | 9991 ± 4.2 | 9984 ± 5.6 | 67.7 ± 5.7 | 0.102 ± 0.034 |
| QuRe [5] | 3.7 ± 1.9 | 6993 ± 1306.3 | 7374 ± 686.5 | 43.8 ± 13.5 | 0.331 ± 0.318 |
| b. Proportion of three samples: 0.5, 0.3, 0.2 (total length of three haplotypes: 30k) |  |  |  |  |  |
| Software | # contigs<br>$\geq 500\text{bp}$ | Longest<br>contig | N50 | Haplotypes<br>coverage % | Error rate % |
| HaploJuice | **3.0 ± 0.0** | **9975 ± 6.3** | **9971 ± 7.8** | **99.7 ± 0.0** | **0.000 ± 0.001** |
| hmmfreq [1] | **3.0 ± 0.0** | 9854 ± 5.8 | 9850 ± 7.6 | 98.5 ± 0.0 | 0.089 ± 0.104 |
| shoRAH [2] | 27.9 ± 6.6 | 9814 ± 118.3 | 9789 ± 113.9 | 97.1 ± 4.7 | 0.591 ± 0.358 |
| SAVAGE [3] | 11.4 ± 3.4 | 9983 ± 8.2 | 436 ± 281.8 | 54.7 ± 7.1 | 0.001 ± 0.005 |
| PredictHaplo [4] | 2.0 ± 0.2 | 9991 ± 3.7 | 9984 ± 5.8 | 68.0 ± 6.6 | 0.087 ± 0.040 |
| QuRe [5] | 4.2 ± 2.2 | 7348 ± 820.8 | 7436 ± 776.9 | 44.9 ± 15.9 | 0.761 ± 0.851 |
| c. Proportion of three samples: 0.6, 0.3, 0.1 (total length of three haplotypes: 30k) |  |  |  |  |  |
| Software | # contigs<br>$\geq 500\text{bp}$ | Longest<br>contig | N50 | Haplotypes<br>coverage % | Error rate % |
| HaploJuice | **3.0 ± 0.0** | **9975 ± 7.3** | **9970 ± 7.7** | **99.7 ± 0.0** | **0.000 ± 0.000** |
| hmmfreq [1] | **3.0 ± 0.0** | 9854 ± 5.6 | 9849 ± 6.2 | 98.5 ± 0.0 | 0.210 ± 0.214 |
| shoRAH [2] | 25.2 ± 5.9 | 9837 ± 115.0 | 9808 ± 113.3 | 97.4 ± 4.8 | 0.749 ± 0.516 |
| SAVAGE [3] | 11.2 ± 3.0 | 9971 ± 20.9 | 419 ± 260.5 | 53.9 ± 6.3 | 0.001 ± 0.006 |
| PredictHaplo [4] | 2.0 ± 0.0 | 9991 ± 3.5 | 9984 ± 4.7 | 66.7 ± 0.0 | 0.089 ± 0.025 |
| QuRe [5] | 3.9 ± 1.9 | 7074 ± 1284.4 | 7300 ± 716.6 | 39.1 ± 14.5 | 0.492 ± 0.597 |
| d. Proportion of three samples: 0.7, 0.2, 0.1 (total length of three haplotypes: 30k) |  |  |  |  |  |
| Software | # contigs<br>$\geq 500\text{bp}$ | Longest<br>contig | N50 | Haplotypes<br>coverage % | Error rate % |
| HaploJuice | **3.0 ± 0.0** | **9976 ± 6.1** | **9971 ± 6.3** | **99.7 ± 0.0** | **0.005 ± 0.048** |
| hmmfreq [1] | **3.0 ± 0.0** | 9855 ± 6.2 | 9850 ± 6.7 | 98.5 ± 0.0 | 0.240 ± 0.220 |
| shoRAH [2] | 20.2 ± 4.7 | 9835 ± 115.0 | 9812 ± 106.4 | 93.8 ± 11.2 | 0.912 ± 0.630 |
| SAVAGE [3] | 15.2 ± 3.0 | 9974 ± 10.6 | 708 ± 161.7 | 65.1 ± 7.0 | 0.001 ± 0.005 |
| PredictHaplo [4] | 2.0 ± 0.0 | 9991 ± 3.8 | 9984 ± 4.7 | 66.7 ± 0.0 | 0.088 ± 0.021 |
| QuRe [5] | 3.6 ± 1.8 | 6787 ± 1333.0 | 7121 ± 809.6 | 28.4 ± 11.2 | 0.319 ± 0.535 |

**Table 3.** Estimated frequencies of three kangaroo sub-samples among the mixture of reads [1] for three amplicons resulted from our method. It revealed the existence of variations on the ratios of the sub-samples when mixing them during the library preparation. 10 data sets were for each amplicon

| Amplicon | Target proportions |  |  | Average estimated proportions (average variation in %) |  |  |
| --- | --- | --- | --- | --- | --- | --- |
| | $f_1$ | $f_2$ | $f_3$ | $f_1$ | $f_2$ | $f_3$ |
| Amplicon 1 | 0.625 | 0.250 | 0.125 | 0.656 (4.9%) | 0.229 (8.3%) | 0.115 (8.0%) |
| Amplicon 2 | 0.625 | 0.250 | 0.125 | 0.640 (2.4%) | 0.246 (1.6%) | 0.114 (8.7%) |
| Amplicon 3 | 0.625 | 0.250 | 0.125 | 0.646 (3.4%) | 0.251 (0.3%) | 0.103 (17.9%) |

**Table 4.** Comparison of performance of different methods on construction of three haplotypes for real kangaroo data sets from the mixture of reads [1] for (a) amplicon 1, (b) amplicon 2, and (c) amplicon 3. There are 10 data sets for each amplicon with total coverage of the reads 1600x. For each data set, the sub-samples were mixed in the proportions: 0.125, 0.25, 0.625. The format of data is: average ± standard deviation. The best value for each column is highlighted among the methods with contigs over 90% coverage on three haplotypes.

| Software | # contigs<br>$\geq 500\text{bp}$ | Longest<br>contig | N50 | Haplotypes<br>coverage % | Error rate % |
| --- | --- | --- | --- | --- | --- |
| HaploJuice | **3.0 ± 0.0** | **4613 ± 2.1** | **4612 ± 2.0** | **99.4 ± 0.0** | **0.05 ± 0.07** |
| hmmfreq [1] | **3.0 ± 0.0** | 4485 ± 0.6 | 4484 ± 0.6 | 96.6 ± 0.0 | 0.26 ± 0.10 |
| shoRAH [2] | 24.0 ± 2.6 | 4592 ± 7.0 | 4592 ± 6.0 | 95.6 ± 10.4 | 1.05 ± 0.32 |
| SAVAGE [3] | 13.2 ± 2.1 | 903 ± 132.3 | 482 ± 169.6 | 47.3 ± 5.2 | 0.02 ± 0.04 |
| PredictHaplo [4] | 1.1 ± 0.3 | 4630 ± 2.0 | 462 ± 1461.3 | 36.5 ± 10.5 | 0.01 ± 0.01 |
| QuRe [5] | 4.0 ± 1.9 | 4343 ± 9.9 | 3909 ± 1373.7 | 74.9 ± 21.8 | 0.42 ± 0.32 |

b. Amplicon 2 (total length of three haplotypes: 12694)
| Software | # contigs<br>$\geq 500\text{bp}$ | Longest<br>contig | N50 | Haplotypes<br>coverage % | Error rate % |
| --- | --- | --- | --- | --- | --- |
| HaploJuice | **3.0 ± 0.0** | **4120 ± 1.5** | **4120 ± 1.5** | **97.4 ± 0.0** | **0.02 ± 0.03** |
| hmmfreq [1] | **3.0 ± 0.0** | 3998 ± 4.0 | 3998 ± 4.0 | 94.5 ± 0.1 | **0.02 ± 0.01** |
| shoRAH [2] | 24.2 ± 5.7 | 4119 ± 14.5 | 4118 ± 12.1 | 90.8 ± 13.5 | 0.41 ± 0.48 |
| SAVAGE [3] | 8.8 ± 3.8 | 1806 ± 761.5 | 572 ± 81.7 | 50.2 ± 4.7 | 0.00 ± 0.00 |
| PredictHaplo [4] | 2.0 ± 0.0 | 4140 ± 2.6 | 4136 ± 0.0 | 65.2 ± 0.0 | 0.00 ± 0.00 |
| QuRe [5] | 2.4 ± 0.7 | 3746 ± 4.7 | 3373 ± 1185.0 | 38.4 ± 14.3 | 0.22 ± 0.28 |

c. Amplicon 3 (total length of three haplotypes: 15391)
| Software | # contigs<br>$\geq 500\text{bp}$ | Longest<br>contig | N50 | Haplotypes<br>coverage % | Error rate % |
| --- | --- | --- | --- | --- | --- |
| HaploJuice | **3.0 ± 0.0** | 5116 ± 9.1 | **5111 ± 7.7** | **99.6 ± 0.1** | **0.01 ± 0.00** |
| hmmfreq [1] | **3.0 ± 0.0** | 5029 ± 3.1 | 5027 ± 3.6 | 98.0 ± 0.1 | 0.23 ± 0.11 |
| shoRAH [2] | 27.6 ± 3.0 | **5132 ± 7.1** | 5111 ± 7.4 | 96.3 ± 10.5 | 1.91 ± 0.44 |
| SAVAGE [3] | 11.8 ± 2.3 | 2510 ± 672 | 550 ± 40.4 | 55.6 ± 4.3 | 0.01 ± 0.01 |
| PredictHaplo [4] | 1.6 ± 0.5 | 5170 ± 3.9 | 3070 ± 2642.4 | 53.3 ± 17.2 | 0.14 ± 0.09 |
| QuRe [5] | 3.0 ± 1.1 | 4567 ± 2.1 | 4106 ± 1442.7 | 35.6 ± 12.5 | 0.25 ± 0.28 |

Apart from the simulated data sets, mixtures of reads from three kangaroo sub-samples [1] were also used to evaluate the performance of the methods. These reads [1] were obtained by short read sequencing of three mitochondrial amplicons on an Illumina platform. The sub-samples were mixed in the proportions: 0.625, 0.25, and 0.125 during the library preparation, and the total coverage of reads is 1600x. There is a total of 30 data sets; 10 data sets for each amplicon (three amplicons in total).

All the reads were aligned against the corresponding amplicon regions on the reference mitochondrial sequence [16] (Genbank accession number NC_027424) by Bowtie 2 [9]. The alignment file is the input of HaploJuice and the estimated sub-sample proportions are listed in Table 3. Although the sub-samples were intentionally mixed in the proportions 0.625, 0.25 and 0.125, variations on the estimated proportions were noticed. For example, for the data sets of amplicon 3, the estimated proportions were 0.646, 0.251, and 0.103 on average. The variation between the estimated proportions and the expected proportions was 6.2% on average, ranging from 0.3% to 17.9%. This revealed the fact that the actual sub-sample proportions in the mixture may be differ from expectation, when the sub-samples are mixed manually during the library preparation.

HaploJuice as well as the other five methods, including Hmmfreq [1], ShoRAH [2], SAVAGE [3], PredictHaplo [4] and QuRe [5], were used to reconstruct the three haplotypes for each amplicon region from the mixture of kangaroo reads. MetaQUAST [15] with default parameters was used to evaluate the resulting contigs against the true haplotypes inferred by deep sequencing [1]. Table 4 shows the summary on the performance of different methods. On average, HaploJuice resulted in contigs with the highest haplotype coverage for all amplicons (97% for amplicon 2 and over 99% for amplicon 1 and 3) among all the methods, and with the lowest (or one of the lowest) error rate among the methods with contigs over 90% haplotype coverage (on average, 0.05% for amplicon 1, 0.02% for amplicon 2, and 0.01% for amplicon 3). Thus, HaploJuice is shown effective from these real data sets.

When comparing the running time between different methods on the Kangaroo data sets, HaploJuice was the fastest, averaging 0.14 minutes for each data set, while other software took from 4 to 139 minutes. The summary is shown in Table 5.

**Table 5.** The average running time (in min) of different methods to reconstruct haplotypes for each Kangaroo data set

| HaploJuice | hmmfreq [1] | ShoRah [2] | SAVAGE [3] | PredictHaplo [4] | QuRe [5] |
| --- | --- | --- | --- | --- | --- |
| 0.14 | 13.53 | 7.81 | 11.21 | 4.30 | 139.93 |

## Discussion

In order to decrease the cost of sequencing, Ranjard, et al. [1] proposed a pooling strategy to mix subsamples in specific known proportions thus simplifying library preparation by removing the need for barcode sequences. According to their experiments on mitochondrial amplicons from three kangaroo sub-samples mixed in proportions 0.625, 0.25, and 0.125, they found that the three haplotypes could be reconstructed effectively using these known frequencies. However, they found that variation of the ratios of sub-samples when mixing due to stochastic experimental effects can decrease the accuracy of haplotype reconstruction. Our research provides an alternative haplotype reconstruction algorithm for Ranjard et al's pooling strategy. We show that estimating the empirical proportions of the mixed sub-samples, prior to the reconstruction the haplotype sequences, significantly increases the accuracy of the approach. As shown from the simulated data sets and the real data sets, our method can, first, accurately identify the underlying sub-sample proportions from a mixture of reads and, second, reconstruct the haplotypes according to these estimated proportions.

The pooling strategy can be applied on a greater number of sequences. Consider a total of n subsamples. A group of three sub-samples of the same species can be mixed in the specific known proportions and applied the same barcode. Thus only 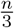 barcodes are required and the cost of the library preparation can be greatly reduced. After sequencing, HaploJuice can be used to assemble the reads associated with the same barcode and reconstruct the three haplotypes for each group of the sub-samples. As shown from the simulated data sets and the real data sets, the high accuracy of assembled haplotypes makes the suggested pooling strategy [1] become more realistic, feasible, and applicable.

Our method relies on aligning reads against a reference sequence. The accuracy of the read alignments affects the effectiveness of our method. In our evaluations, we only used alignments reported by Bowtie 2 [9] with mapping quality of at least 20. Whereas we understand that coverage varies along the haplotype, but we assume that ratios of the read coverage for each haplotype at each location follows the same multinomial distribution. If a region on some haplotypes is very different from the reference sequence, reads from this region may not align to the reference, and the induced read coverage for those haplotypes may decrease substantially. The bias in the induced read coverage ratio can cause misleading results, because of its deviation from the common multinomial distribution. Therefore, this method is designed for the pooling strategy applied on the sub-samples that align well with the reference sequence.

HaploJuice assumes that the number of haplotypes is known in advance. There is no equivalent assumption with ShoRAH [2], SAVAGE [3], PredictHaplo [4] and QuRe [5]. Nonetheless, these are the only available software for haplotype reconstruction from a pool of reads originating from a mixture of different sub-samples. We expect that the effectiveness of haplotype reconstruction using these methods are also likely to be improved if the number of haplotypes is known in advance. One reasonable approach to assemble the reads from a sample with unknown number of haplotypes is therefore to develop a statistical method to estimate the number of haplotypes from a mixture of reads, and then reconstruct the haplotypes using our method according to this estimated number of haplotypes.

## Conclusion

HaploJuice is designed for the reconstruction of three pooled haplotypes from a mixture of short sequencing reads obtained under the strategy proposed by Ranjard, et al. [1]. As shown from the simulated data sets and the real data sets, HaploJuice provides high accuracy in haplotype reconstruction, thus increasing the estimation efficiency of Ranjard et al.’s pooling strategy.

## Methods

### Estimation of sample proportions

HaploJuice requires an alignment of short-read sequences against a reference sequence. All reads are aligned to the reference sequence using Bowtie 2 [9]. Only the reads which are aligned at unique positions on the reference are considered. The alignment of each read has a starting and an ending position on the reference. A sliding window approach is used.

Let *W* be the set of overlapping windows. For each window *w* ∈ *W*, we collect the reads that are aligned across the whole window. We extract the corresponding sub-sequences according to the window’s bounds, and obtain the set of unique sub-sequences *T_w_* = {*t*_*w*1_, *t_w_*_2_, …} and the frequencies *G_w_* = {*g_w_*_1_, *g_w_*_2_, …} where *g_wi_* is the number of reads with subsequence *t_wi_*. The sub-sequences inside *T_w_* are sorted in decreasing order of frequencies.

Say *n* sub-samples are pooled with unknown proportions *f*_1_,*f*_2_,…,*f*_n_ where *f*_1_ > *f*_2_ > … > *f*_n_. When there is no sequencing error and each sub-sample is from a unique haploid sequence, each subsample should produce only one subsequence in *T_w_*. In those regions where two or more sub-samples are identical, the sub-sequences originating from these sub-samples will be the same. For each sliding window, the number of possible combinations of *n* samples producing sub-sequences, i.e. the number of possible partitions of a set with n different elements (where each element represents a sub-sample, and the elements in the same partition are regarded as the sub-samples producing the same sub-sequences), is the Bell number *B_n_* [17]. Each case will lead to different expected frequencies of the sub-sequences.

However, under real sequencing conditions, the number of sub-sequences in each window may be greater than *n*, because some erroneous sub-sequences are created by sequencing errors. We assume that the frequencies of erroneous sub-sequences are always lower than that of real sub-sequences. For each window, we only consider the top-n most frequent sub-sequences. Table 6 lists the expected frequencies of the sub-sequences for all cases when *n* = 3.

Let *p_ki_* be the *i*-th expected frequency for case *k*. Assume the observed frequencies of the subsequences in a window *w* ∈ *W* follow a multinomial distribution. The likelihood value for the window *w*, (*L*(*w*)),is computed as follows:

**Table 6.** The expected frequencies of top-n most frequent sub-sequences for a mixture from 3 samples. This is a total of *B*_3_ = 5 cases. *f_e_* and *f_e’_* are the proportions of erroneous sequences.

| Case | Expected frequencies of sub-sequences |  |  |
| --- | --- | --- | --- |
| 1 | $f_1$ | $f_2$ | $f_3$ |
| 2 | $f_1 + f_2$ | $f_3$ | $f_e$ |
| 3 | $f_1 + f_3$ | $f_2$ | $f_e$ |
| 4 | $f_2 + f_3$ | $f_1$ | $f_e$ |
| 5 | $f_1 + f_2 + f_3$ | $f_e$ | $f_{e'}$ |

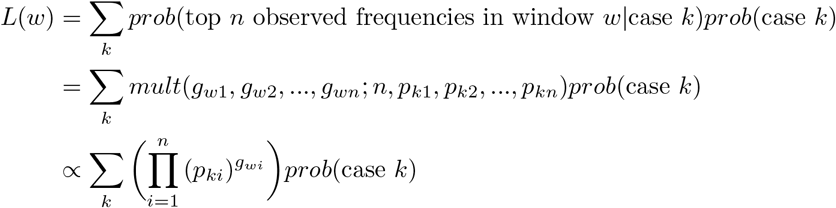

The probability of the case *k* (i.e. *prob*(case *k*)) is estimated by the following equation:

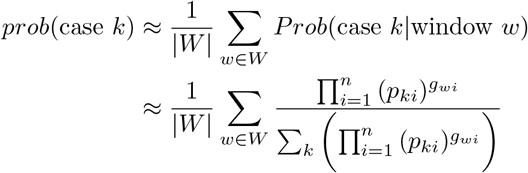

And the overall log-likelihood value (*logL*) for all the windows *w* ∈ *W* is:

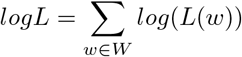

The optimal values of *f*_1_,*f*_2_,…,*f_n_*,*f_e_*,*f_e’_* are computed such that the overall log-likelihood value (*logL*) is maximum. In practice, the following constraints are used: *f*_1_ ≥ *f*_2_ ≥ … *f_n_* ≥ *f_e_* ≥ *f_e’_* and *f_e_* ≤ *b*, where *b* is an upper limit for the frequency of an erroneous subsequence. The estimated sample proportions are the optimal values of *f*_1_,*f*_2_,…,*f_n_*.

## Reconstruction of haplotype sequences

The next step is to reconstruct the haplotype sequences according to the sub-sample proportions estimated in the previous step. We assume that each sub-sample is generated from a unique haploid sequence (i.e. haplotype). If we can identify the corresponding sub-sequence of each haplotype for every sliding window, then the haplotype sequences can be reconstructed by combining the sub-sequences from all the windows. However, in practice, it is not obvious, because the real sub-sequences are usually mixed with erroneous sub-sequences caused by sequencing errors. Moreover, multiple haplotypes may share the same sub-sequence and the observed frequencies of the sub-sequences may deviate from expectation at some positions.

A dynamic programming approach was used to reconstruct multiple haplotype sequences simultaneously, by considering all the cases for each window, and choosing the best arrangement with the maximum likelihood value.

Consider a sliding window *w* ∈ *W* and the top-*n* most frequent sub-sequences (i.e. *t_w_*_1_,*t_w_*_2_,…,*t_wn_*) in the window. Since each haplotype can generate one sub-sequence, there are *n^n^* possible cases to generate *n* different sub-sequences by *n* haplotypes (considering that multiple haplotypes can generate the same sub-sequence and some sub-sequences can be erroneous), and each case will lead to a different set of expected frequencies of the sub-sequences. Table 7 lists all 27 possible cases and the expected frequencies of the sub-sequences when *n* = 3.

**Table 7.** There are a total of 27 cases for generating 3 sub-sequences by 3 haplotypes. *hi* represents that the sub-sequence is generated from haplotype *i*, and ‘erroneous’ represents the erroneous sub-sequences. *fi* is the estimated proportion of sample *i*, and *f_e_*, *f_e’_* are the proportions of erroneous sub-sequences.

| Case | Haplotypes which generate the sub-sequences |  |  | Expected frequencies |  |  |
| --- | --- | --- | --- | --- | --- | --- |
|  | subseq1 | subseq2 | subseq3 | subseq1 | subseq2 | subseq3 |
| 1 | $h_1$ | $h_2$ | $h_3$ | $f_1$ | $f_2$ | $f_3$ |
| 2 | $h_1$ | $h_3$ | $h_2$ | $f_1$ | $f_3$ | $f_2$ |
| 3 | $h_2$ | $h_1$ | $h_3$ | $f_2$ | $f_1$ | $f_3$ |
| 4 | $h_2$ | $h_3$ | $h_1$ | $f_2$ | $f_3$ | $f_1$ |
| 5 | $h_3$ | $h_1$ | $h_2$ | $f_3$ | $f_1$ | $f_2$ |
| 6 | $h_3$ | $h_2$ | $h_1$ | $f_3$ | $f_2$ | $f_1$ |
| 7 | $h_1$ & $h_2$ | $h_3$ | erroneous | $f_1 + f_2$ | $f_3$ | $f_e$ |
| 8 | $h_3$ | $h_1$ & $h_2$ | erroneous | $f_3$ | $f_1 + f_2$ | $f_e$ |
| ... | ... | ... | ... | ... | ... | ... |
| 26 | erroneous | $h_1$ & $h_2$ & $h_3$ | erroneous | $f_e$ | $f_1 + f_2 + f_3$ | $f_{e'}$ |
| 27 | erroneous | erroneous | $h_1$ & $h_2$ & $h_3$ | $f_e$ | $f_{e'}$ | $f_1 + f_2 + f_3$ |

Define *A*(*w*, *k*) = (*t*_1_, …, *t_n_*) as an assignment of the haplotypes to the sub-sequences in sliding window *w* when case *k* is considered (i.e. *i*-th haplotype generates sub-sequence *t_i_*). For example, as shown in Table 7, for *n* = 3 and case 7, *A*(*w*, 7) = (*t_w_*_1_,*t_w_*_1_, *t_w_*_2_) (i.e. the observed sub-sequence with the highest frequency in window w is generated from both the first and the second haplotypes, while the observed alignment with the second highest frequency is generated from the third haplotype).

Define δ(*A*(*w*, *k*), *A*(*w’, k’*)) as the compatibility between two assignments *A*(*w*, *k*) = (*t*_1_, …, *t_n_*) and *A*(*w’*,*k’*) = (*t’*_1_, …, *t’_n_*) and δ(*A*(*w*, *k*), *A*(*w’, k’*)) = 1 if, for all 1 ≤ *i* ≤ *n*, two sub-sequences *t_i_* and *t’_i_* are exactly the same in their overlapped region. Mathematically, if the window size is *d*, the two windows overlap *l* bases, and window *w* is before window *w’*,

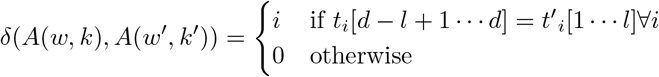

We begin from a starting window *w_s_* ∈ *W* and consider all possible *n^n^* assignments in *w_s_*. Then we consider the left and the right windows besides *w_s_*, and continue until all the windows have been considered. The optimal construction of *n* haplotypes is the set of compatible assignments for all the windows with the maximum log-likelihood value. The following dynamic programming approach is used to compute the optimal compatible assignments for all the windows.

Given a starting window *w_s_* ∈ *W*, define ζ(*k_s_,k_t_,w_t_*), where *w*_*t*_ ∈ *W*,1 ≤ *k_s_*,*k_t_* ≤ *n^n^*, as the maximum log-likelihood value of the optimal compatible assignments for the consecutive windows from *w*_*s*_ to *w*_*t*_ with assignment *A*(*w_s_*, *k_s_*) in window *w_s_* and assignment *A*(*w_t_*, *k_t_*) in window *w_t_*. If *s* < *t*, the assignment is proceeded from left to right, while if *t* < *s*, the assignment is proceeded from right to left. Without loss of generality, considering the situation that the haplotype assignment is proceeded from left to right, the recursive formula of ζ(*k_s_, k_t_, w_t_*) is defined as:

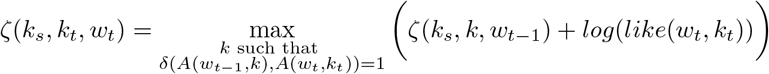

where *like*(*w_t_*, *k_t_*) is the likelihood value of the observed frequencies of the sub-sequences in window *w_t_* when assignment *A*(*w_t_*, *k_t_*) is selected.

Let *q_ki_* be the *i*-th largest expected frequency for case *k*.

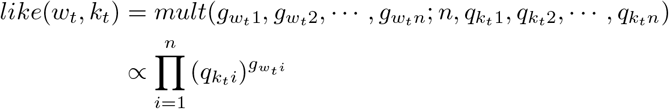

Therefore,

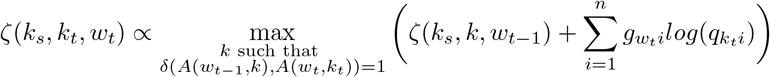

In order to increase the accuracy of the haplotype construction, we reconstruct the haplotypes starting from a relatively reliable window 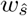 with much dissimilarity between the haplotypes. When *n* = 3, we locate the window 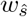 which have the greatest value of likelihood value for the case when each haplotype is assigned to different sub-sequence. Let the first and the last window on the haplotype region be *w_1_* and *w_1ast_*. The haplotypes are constructed in both directions from the window 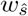 to the beginning and to the ending of the haplotypes, respectively Considering the different case 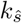 for the starting window 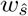
, the log-likelihood value of the optimal set of compatible assignments for the whole haplotype region is:

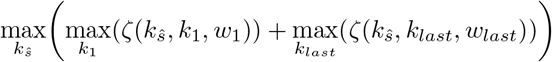

## Availability of data and materials

The software HaploJuice and the simulated datasets are available in OSF repository: https://osf.io/b8nmf/(DOI:10.17605/OSF.IO/B8NMF)

